# Assessing relationships between chromatin interactions and regulatory genomic activities using the self-organizing map

**DOI:** 10.1101/2020.02.20.951616

**Authors:** Timothy Kunz, Lila Rieber, Shaun Mahony

## Abstract

Few existing methods enable the visualization of relationships between regulatory genomic activities and genome organization as captured by Hi-C experimental data. Genome-wide Hi-C datasets are often displayed using “heatmap” matrices, but it is difficult to intuit from these heatmaps which biochemical activities are compartmentalized together. High-dimensional Hi-C data vectors can alternatively be projected onto three-dimensional space using dimensionality reduction techniques. The resulting three-dimensional structures can serve as scaffolds for projecting other forms of genomic information, thereby enabling the exploration of relationships between genome organization and various genome annotations. However, while three-dimensional models are contextually appropriate for chromatin interaction data, some analyses and visualizations may be more intuitively and conveniently performed in two-dimensional space.

We present a novel approach to the visualization and analysis of chromatin organization based on the Self-Organizing Map (SOM). The SOM algorithm provides a two-dimensional manifold which adapts to represent the high dimensional chromatin interaction space. The resulting data structure can then be used to assess the relationships between regulatory genomic activities and chromatin interactions. For example, given a set of genomic coordinates corresponding to a given biochemical activity, the degree to which this activity is segregated or compartmentalized in chromatin interaction space can be intuitively visualized on the 2D SOM grid and quantified using Lorenz curve analysis. We demonstrate our approach for exploratory analysis of genome compartmentalization in a high-resolution Hi-C dataset from the human GM12878 cell line. Our SOM-based approach provides an intuitive visualization of the large-scale structure of Hi-C data and serves as a platform for integrative analyses of the relationships between various genomic activities and genome organization.

## INTRODUCTION

The Hi-C assay captures pairwise interactions between loci across the entire genome [1]. The procedure begins with the isolation of intact nuclei and crosslinking via formaldehyde. Crosslinked chromatin is then fragmented, and the ends of the resulting fragments are biotin labeled. A random ligation step joins the ends of DNA fragments that are in close physical proximity, typically because they are crosslinked in the same complex. After reversing crosslinks, purifying DNA, and further DNA shearing, the fragments that contain ligation products are immunopurified via the biotin tag. The resulting DNA fragments will contain instances where the two ends of the same DNA molecule were ligated together (“self-ligations”) and instances where longer range interactions resulted in intermolecular ligations. Fragments are subjected to paired-end sequencing, and aligned to the genome, resulting in data representing tens to hundreds of millions of pairwise interaction “contacts”.

Hi-C data is processed by binning the genome (where the bin size is dependent on sequencing depth and molecular complexity) and counting the interaction contacts between each pair of bins. The contact frequencies recorded in the resulting interaction matrix are inversely proportional to the average 3D distance between loci in the cell population [1]. Interaction matrices are typically visualized using a heatmap (**Figure 1a**). While the matrices are visually dominated by the products of self-ligations along the matrix diagonal, non-uniform interaction frequencies between loci can also be seen off-diagonal. The patterns of preferential interactions can be more clearly visualized by normalizing the observed interaction frequencies using frequencies expected at each linear genomic distance, which produces an observed/expected (O/E) matrix (**Figure 1b**).

**Figure 1:**
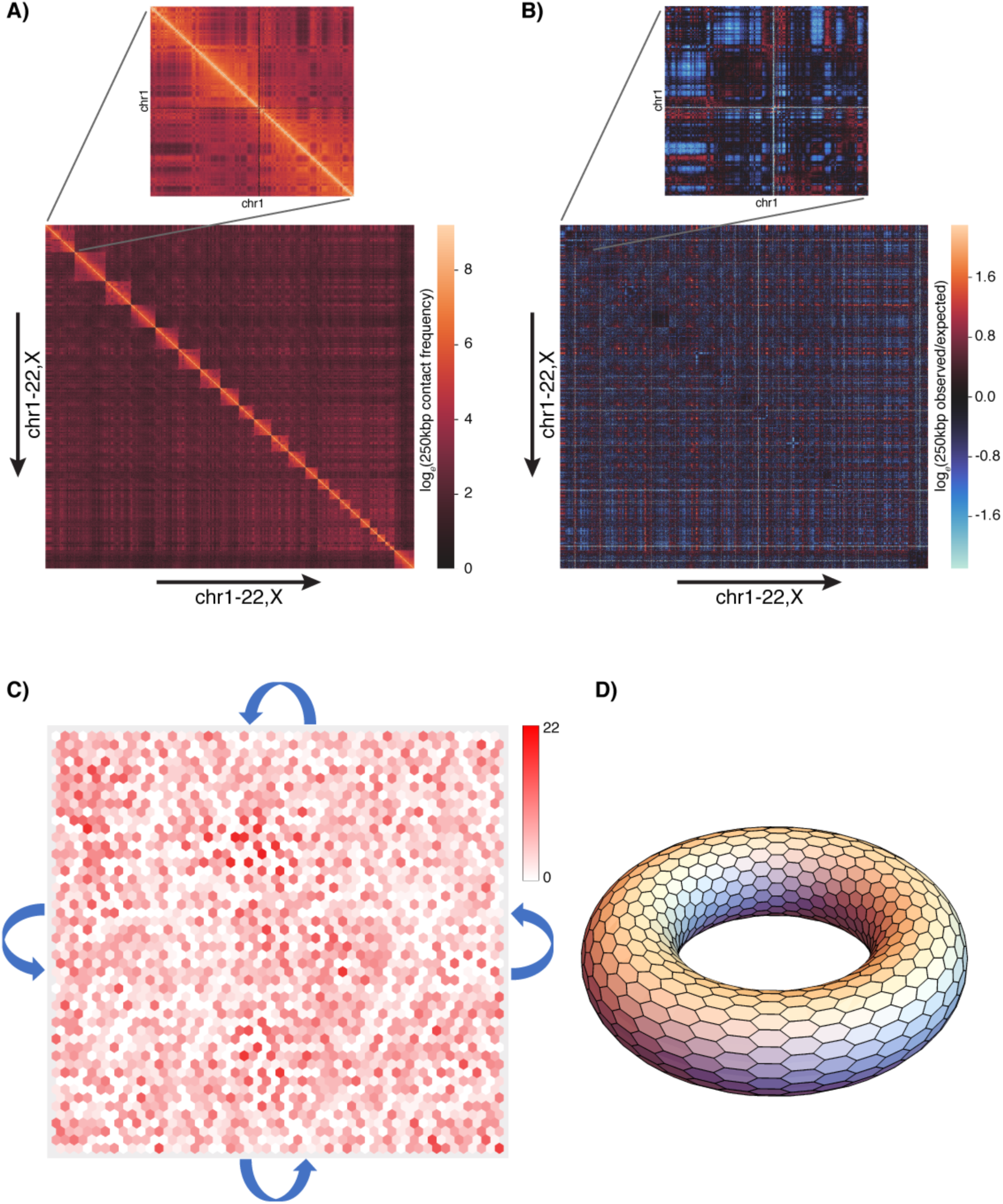
Overview of ChromoSOM, a Self-Organizing Map for visualizing chromatin organization. **A)** Chromatin interaction frequencies in 250kbp intervals are calculated genome-wide from GM12878 Hi-C data, and **B)** transformed into normalized observed/expected values. **C)** A 50×50 hexagonal grid Self-Organizing Map is trained using the rows of the observed/expected matrix. Training datapoints are distributed somewhat uniformly over the trained lattice. **D)** The lattice is defined to have a toroidal structure, in which the opposing edges and corners are connected.

In the interaction and O/E matrices, each row represents an *N*-dimensional interaction vector for a given bin on the genome. Several dimensionality reduction approaches have been applied to reduce the complexity of these high-dimensional vectors. For example, Principal Component Analysis (PCA) is often applied to a correlation matrix derived from the O/E matrix [1,2]. The first principal component typically corresponds to a broad division between two major compartments within the nucleus, one containing active genomic processes (compartment A) and one containing repressed chromatin (compartment B). PCA-derived compartment analysis can thus be thought of as reducing the complexity of Hi-C data onto a single dimension. We and others have also generated methods that convert Hi-C interaction matrices into 3-dimensional structures, which represent the average conformation of chromosomes in a given cell population [3–13]. While the methodologies vary from modeling-based approaches using Markov Chain Monte Carlo [3–7] to optimization-based approaches using multidimensional scaling [8–13], the effect of all such approaches is an embedding of the *N*-dimensional interaction information into 3D space.

Once chromatin interactions have been characterized in a given cell type, it is natural to ask whether they are associated with regulatory activities such as transcription, chromatin accessibility, histone modifications, and protein-DNA interactions [1]. However, few existing approaches enable an intuitive visualization and quantification of relationships between chromatin interactions and regulatory signals. The locations of regulatory signals can be correlated with compartment annotations, but this may miss more subtle relationships between regulatory activities and sub-compartment level chromatin interactions. Regulatory signals can also be painted onto a 3D genome structure [9,13,14], analogous to how signal tracks can be presented along the linear genome in a 1D genome browser. While genome structures can provide an intuitive framework for visualizing the 3D context of specific regulatory activities, it is difficult to visualize overall trends as the entire 3D structure cannot be seen in a single static image. Likewise, it is also difficult to quantify the overall associations between a given regulatory activity and the 3D structure.

Here, we present a new approach to visualizing and quantifying relationships between chromatin interaction space and regulatory genomic activities using the Self-Organizing Map (SOM) [15,16]. The SOM is a popular machine learning approach to non-linear dimensionality reduction. The SOM’s training procedure iteratively fits a 2-dimensional output lattice of nodes (or “neurons”) to the *N*-dimensional input space. The nodes encapsulate *N*-dimensional feature vectors that adapt to represent some component of the input space. However, relationships between neighboring nodes on the 2D lattice are constrained such that the output lattice preserves the topology of the input space. The SOM has been extensively used in biological applications, for example to provide dimensionality reduction of high-dimensional gene expression patterns [17,18], regulatory DNA motif features [19,20], genomic and metagenomic sequence *k*-mer frequency profiles [21–25], and regulatory genomic signal profiles [26].

In our application to the analysis of chromatin interaction vectors, we define each SOM node as containing a feature vector of the same dimensionality as chromatin interaction vectors in the training set. During training, each node adapts to represent some set of similar training vectors, in effect clustering the underlying genomic loci “within” the node. Because of the SOM’s topology preserving properties, nearby nodes on the SOM’s 2D lattice will end up representing similar chromatin interaction vectors, and thus will contain sets of genomic loci that are nearby each other in the nucleus. Therefore, SOM training should have the effect of projecting genomic loci onto the 2D SOM lattice, where the distribution of loci within that lattice should reflect their relative spatial organization within the nucleus.

We demonstrate that SOMs trained with chromatin interaction vectors have two advantages for characterizing the relationships between chromatin organization and regulatory activities. Firstly, the trained SOM’s output lattice is an easy-to-visualize 2D grid that represents the distribution of genomic loci in chromatin interaction space. The locations of regulatory events (e.g. protein-DNA binding events or histone modifications) can be highlighted on that grid, thus visualizing the distribution of the regulatory activity with respect to chromatin interaction space. Secondly, because each of the SOM’s output lattice nodes represent a fixed set of genomic loci, we can easily count the number of regulatory events that are “clustered” in each node. We show that a modified Lorenz curve analysis can be used to quantify the non-uniformity of a given regulatory activity over the nodes. Since the lattice represents chromatin interaction space, a non-uniform distribution of regulatory events on the SOM can be interpreted as an association between the regulatory activity and some aspect of genome organization.

As a proof of principle, we demonstrate our approach by training a SOM using Hi-C interaction data from the GM12878 cell line. This SOM is then used to assess the distribution of numerous chromatin activities, including histone modifications, chromatin accessibility, and protein-DNA interactions.

## METHODS

### Data

Hi-C data for GM12878 was sourced from [27]. Intra-chromosomal and inter-chromosomal contact frequency matrices from this study were downloaded from the GEO archive (accession GSE63525). While the contact frequency matrices are provided at several resolutions, we chose to focus on a bin size of 250kbp so that inter-chromosomal interaction frequencies were not too sparse. Contact frequency matrices were normalized using Knight & Ruiz matrix balancing factors [28], also downloaded from GSE63525. Rows/columns with zero interactions were removed. Normalized contact frequencies were then converted into observed/expected values, where expected values are calculated as the average interaction frequencies observed at a given genomic distance (calculated separately per chromosome) for intrachromosomal interactions, and the average interaction frequency between all bins in a given pair of chromosomes for interchromosomal interactions. Finally, a single whole genome interaction matrix was constructed using log-transformed observed/expected values. Since the original contact frequency matrices were generated using Hi-C data that was mapped to hg19, all data presented in this study was mapped to that version of the genome.

Regulatory activities were sourced from the ENCODE project portal [29] (https://www.encodeproject.org). We downloaded narrowPeak BED files for all available histone ChIP-seq, TF ChIP-seq, and DNase-seq experiments in GM12878 (hg19). We removed ChIP-seq datasets that contained fewer than 1,000 peaks, and also removed one DNase-seq dataset that contained over 400,000 peaks. This left 3 DNase-seq datasets, 13 histone modification ChIP-seq datasets, and 150 transcription factor ChIP-seq datasets. Finally, we downloaded IDEAS genome segmentation results for 127 human cell types [30], and extracted the annotations corresponding to GM12878.

### Self-Organizing Map

Our SOM implementation, named ChromoSOM, defines the output lattice as a grid of hexagonal nodes (**Figure 1c**). While visualized as a 2D grid, the structure of the output lattice is defined to be toroidal (i.e. opposing edges and all corners are defined to be adjacent) (**Figure 1d**). Each node contains an *N*-dimensional weight vector, where *N* is the number of bins on the genome (i.e. the number of columns in the chromatin interaction matrix). Weight vectors are initialized as being equal to a randomly chosen data point.

Training proceeds using the batch SOM training algorithm. Each training iteration consists of an assignment step and an update step. During the assignment step, each training data point is assigned to the SOM node whose weight vector is most similar according to Pearson correlation. During the update step, the weight vector of each node is updated to reflect the assignment of the data points. Every data point is considered during the updating of each node. The weight vectors of each node are updated by:

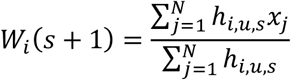

Where: *s* is the current iteration; *W*_*i*_ is the weight vector of node *i*; *N* is the number of data points in the set; *x*_*j*_ is the data vector of data point *j*; and *h*_*i,u,s*_ is the neighborhood function. The neighborhood function is in turn defined as follows:

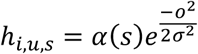

Where: *u* is the SOM node that contains the data point *j* under consideration; *o* is the hexagonal distance on the SOM grid between nodes *i* and *u*; *σ* is the variance of the Gaussian kernel for the current iteration; and *α*(*s*) is the learning rate of the current for the current iteration.

The learning rate shrinks linearly during training from 1.0 to 0.01. The variance parameter also shrinks linearly from 1.2 to 0.2. The SOM is trained for 1,000 iterations. Training is repeated with random initializations 10 times. We save the SOM that has the highest quality, where quality is defined as the average similarity between data points and their assigned nodes [26].

### Analyzing sets of genomic loci using a trained SOM

At the end of the training process, each training point is associated with its best matching node. Since each training data point represents the chromatin interaction profile for a given 250kbp genomic locus, we can think of the nodes as containing a set of zero to many genomic loci. We can thus easily map any set of genomic coordinates to the trained SOM by assigning them to the node that contains an overlapping training data point locus. Mapping a given regulatory activity to the SOM thus consists of taking all loci displaying that activity (e.g. a set of ChIP-seq peaks) and finding the frequencies that they overlap the loci in each SOM node.

To compare the SOM mapping distributions of two genomic activities, we treat the relative mapping frequencies of each dataset to the SOM as a pair of 1D vectors, and perform Pearson correlation between them. To assess the degree to which a given genomic activity is non-uniformly distributed over the SOM, we first order the SOM nodes from lowest to highest overlap with the coordinates defining the activity. We then produce a Lorenz curve [31] (which we term the observed Lorenz curve) that defines the cumulative fraction of query coordinates that are assigned to a cumulative fraction of SOM nodes. Next, we generate 1,000 randomly sampled (with replacement) sets of training data points, where each set contains the same number of training data points as the number of coordinates in the genomic activity under examination. For each set of random training points, we build a Lorenz curve using their node assignments as defined during the SOM training process. The average Lorenz curve over this set of 1,000 is termed the comparison curve. This comparison curve accounts for non-uniformity in the SOM assignment distribution that is merely due to some nodes containing more data points than others at the end of training. We then calculate the area under the observed Lorenz curve (B), and the difference between the area under the comparison curve and the observed curve (A). Our modified Gini coefficient is defined as the ratio: A/(A+B).

## RESULTS

### Self-Organizing Maps can encapsulate chromatin interaction information

We trained a Self-Organizing Map using genome-wide observed/expected Hi-C interaction vectors from the GM12878 human cell line (**Figure 1B**) [27]. A bin size of 250kbp was chosen to enable quantification of inter-chromosomal interactions with sufficient coverage. The SOM’s output lattice was chosen to be a 50×50 hexagonal grid with a toroidal topology, which has been shown to improve the stability of SOM training [16] (**Figure 1C,D**). Since each Hi-C interaction vector represents a 250kbp genomic locus, the SOM training procedure has the effect of clustering zero to many loci in each node on the output lattice. As can be seen in **Figure 1C**, our training procedure results in a relatively smooth distribution of genomic loci across the SOM nodes.

In order to demonstrate that the SOM has appropriately encapsulated chromatin interaction information, we compare the loci clustering represented by the SOM output lattice with previous chromatin compartment annotations produced using the same dataset [27] (**Figure 2A**). Specifically, we map loci that were annotated as occurring in compartment A (active) or compartment B (repressed) to SOM nodes that contain overlapping genomic bins. As shown in **Figure 2A**, compartments A and B loci are well-separated on the 2D SOM lattice. We further map the locations of a finer-grained 6-level sub-compartment annotation [27] to the SOM, and again find that these loci are separated from one another on the SOM lattice. These results suggest that the arrangement of the SOM nodes encodes aspects of chromatin organization within the nucleus.

**Figure 2:**
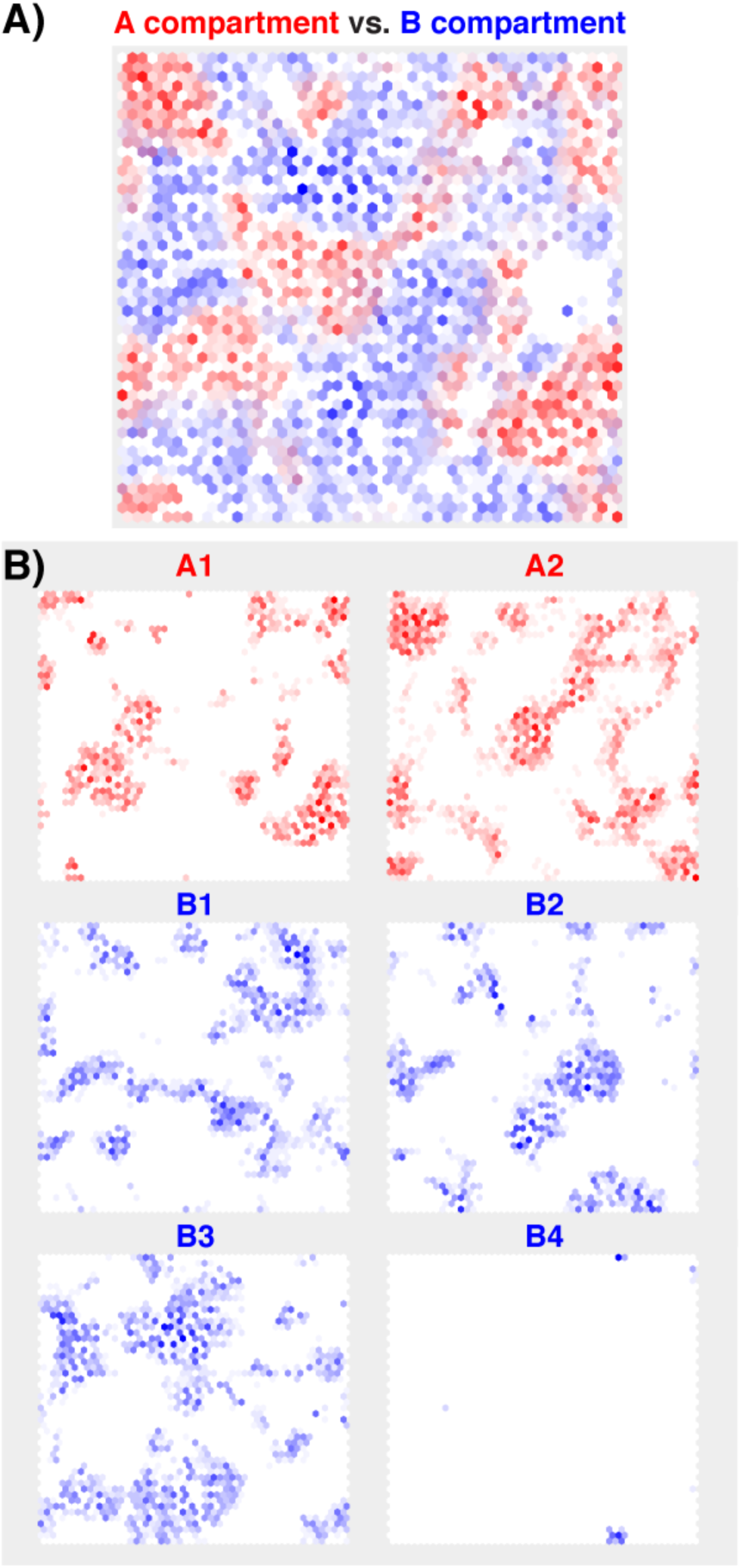
**A)** Comparison of the locations of compartment A (red) and compartment B (blue) loci on the trained SOM. **B)** Projections of the locations of six previously defined sub-compartments on the trained SOM [27].

We further mapped the locations of SOM nodes that contain each individual chromosome’s loci (**Figure 3**). Each chromosome’s loci are somewhat separated from other chromosomes on the SOM lattice, although most chromosomes are mapped in several disjointed clusters. The separation between chromosomes may reflect the existence of distinct chromosome territories within the nucleus [32], while the fragmentation into several clusters per chromosome reflects degrees of (sub-)compartmentalization.

**Figure 3:**
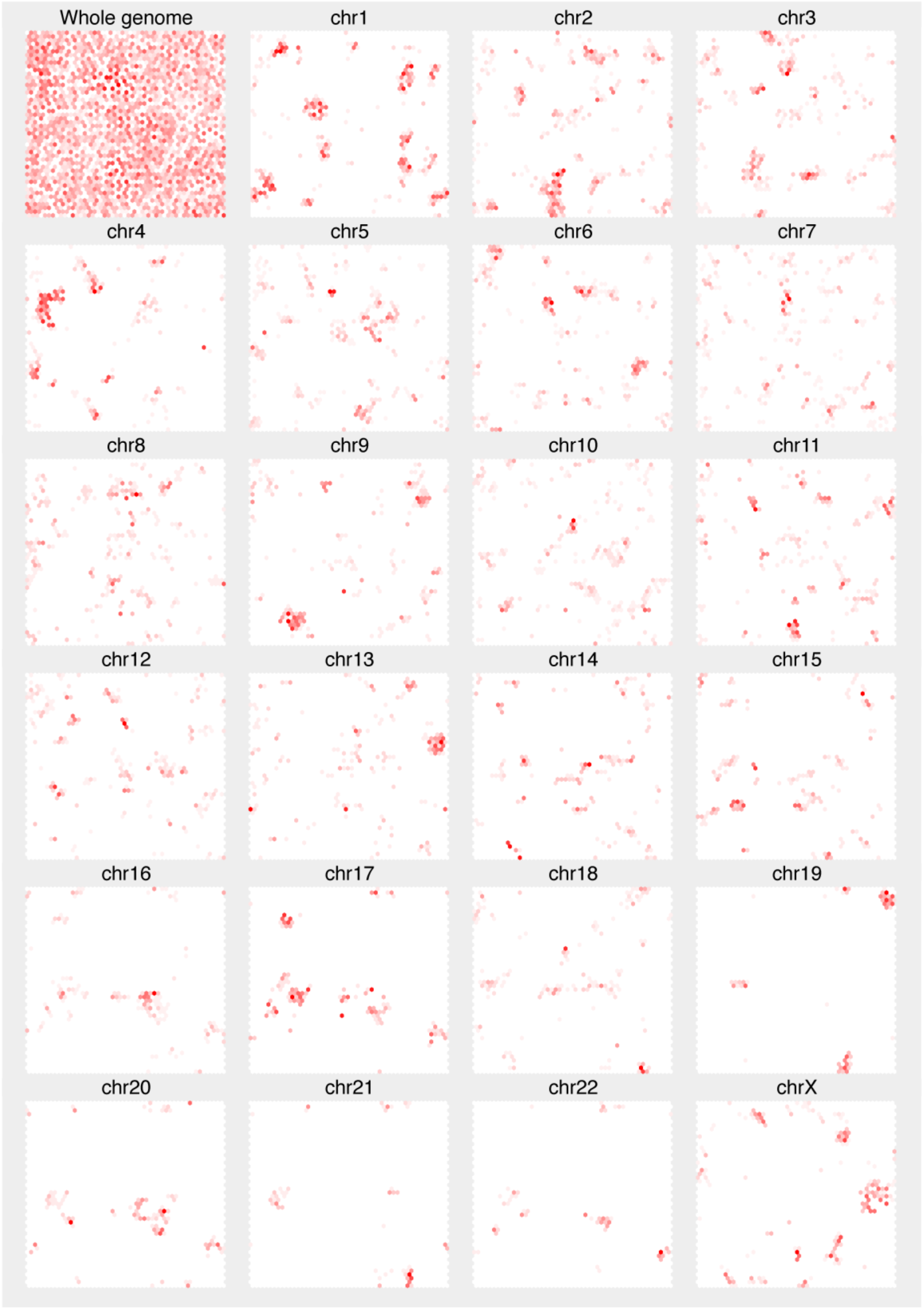
Projection of loci from each individual chromosome on the trained SOM.

### The SOM enables visualization of the spatial distribution of genomic regulatory activities

As demonstrated above, the organization of nodes in a trained SOM encapsulates aspects of genomic spatial organization within the nucleus. Any genomic activity that can be mapped to the genome can be projected onto a trained SOM by assigning loci displaying the activity to nodes that encapsulate overlapping training loci. Doing so implicitly allows us to assess the spatial distribution of the genomic activity within the nucleus, but visualized on the two-dimensional manifold formed by the SOM’s output lattice.

We demonstrate our approach by mapping 16 DNase-seq and histone modification ChIP-seq experiments performed in GM12878 cells to the SOM trained using GM12878 Hi-C data (**Figure 4**). It is apparent from **Figure 4** that the distribution of several histone modification ChIP-seq signals on the SOM is visually similar to the distribution of A compartment loci. For example, histone modifications associated with transcriptional elongation (H3K36me3), transcriptional initiation (H3K4me3), and enhancer activities (H3K4me1 & H3K27ac) all map to the SOM in a manner similar to A compartment annotations. These similar SOM enrichment patterns are unsurprising, as transcription and active regulatory processes are expected to be enriched in the A compartment [1]. However, these comparisons suggest an approach for assessing whether sets of genomic activities are correlated in their spatial distribution within the nucleus.

**Figure 4:**
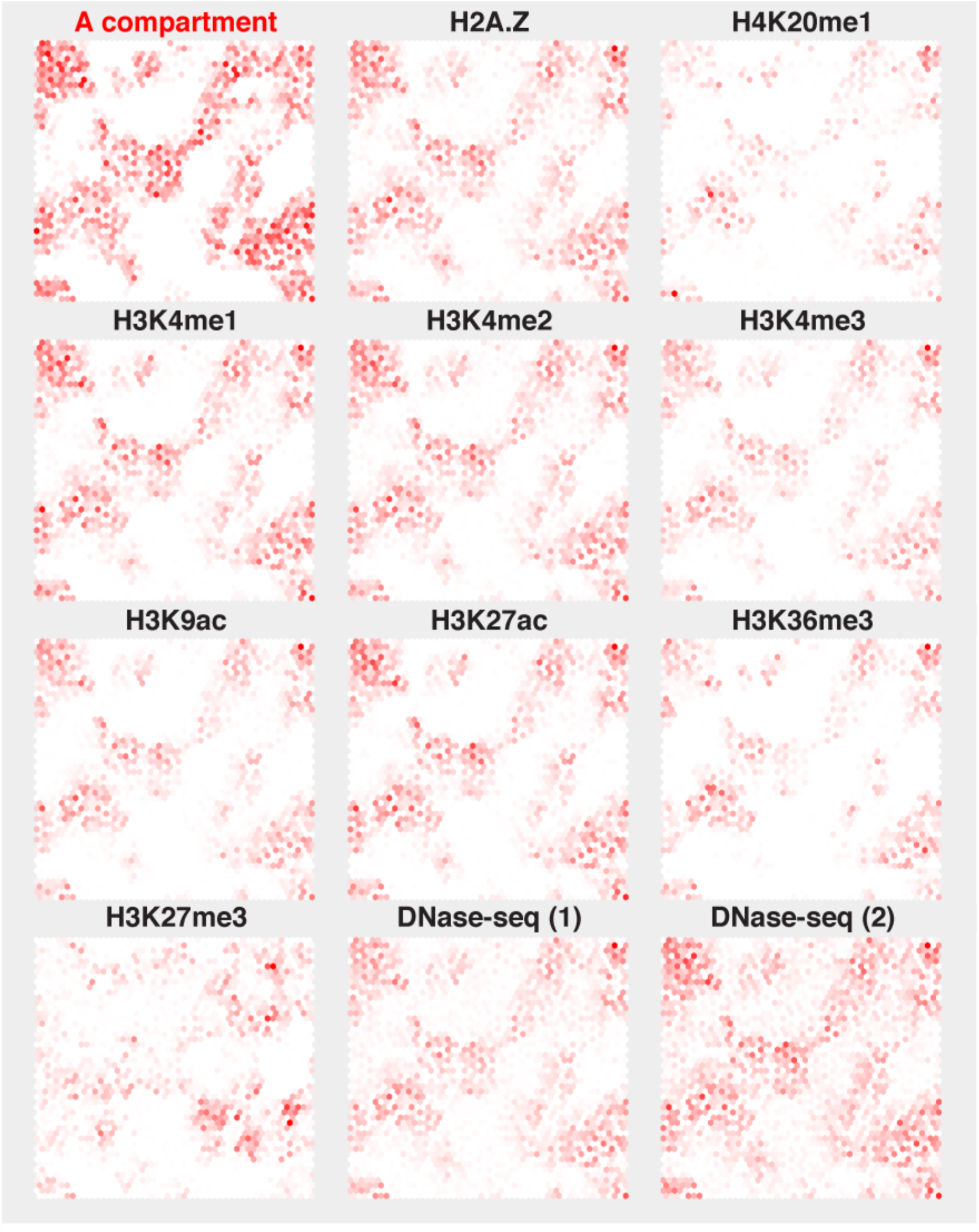
SOM projections of peaks from a selection of 11 DNase-seq and histone mark ChIP-seq experiments performed in the GM12878 cell line.

We can quantify the degree to which sets of genomic annotations or activities are similarly spatially distributed by taking advantage of the discretized nature of the SOM lattice. A dataset’s enrichment pattern over the SOM’s nodes can be treated as a vector, and thus the relationship between two dataset’s enrichment patterns can be assessed by correlation. Comparing the distributions of DNase-seq and histone modification ChIP-seq datasets to A and B compartments confirms that the spatial distributions of most assessed histone modifications are highly correlated with that of the A compartment, and lowly or negatively correlated with that of the B compartment (**Table 1**). An exception is H3K27me3, a histone modification associated with Polycomb repression, which is positively correlated with both A and B compartments.

**Table 1:** Correlations between SOM projections of selected DNase-seq and histone mark ChIP-seq datasets and the SOM projections of compartment A & B loci.

| Dataset | SOM corr. with<br>A compartment | SOM corr. with<br>B compartment |
| --- | --- | --- |
| H3K4me1 ENCFF921LKB | 0.85 | -0.06 |
| H3K27ac ENCFF816AHV | 0.82 | -0.08 |
| H3K4me2 ENCFF983SMS | 0.86 | -0.03 |
| H3K4me3 ENCFF795URC | 0.82 | -0.06 |
| H3K9ac ENCFF052MHA | 0.78 | -0.10 |
| H3K4me3 ENCFF295GNH | 0.81 | -0.07 |
| H3K79me2 ENCFF357HZM | 0.77 | -0.11 |
| H2A.Z ENCFF584NAD | 0.85 | 0.00 |
| H3K4me3 ENCFF354MGT | 0.78 | -0.07 |
| DNase-seq ENCFF097LEF | 0.85 | 0.04 |
| H3K36me3 ENCFF479XLN | 0.68 | -0.12 |
| DNase-seq ENCFF273MVV | 0.87 | 0.13 |
| DNase-seq ENCFF804BNU | 0.71 | 0.03 |
| H4K20me1 ENCFF308WNH | 0.58 | -0.07 |
| H3K27me3 ENCFF851UKZ | 0.43 | 0.20 |
| H3K27me3 ENCFF247VUO | 0.47 | 0.27 |

### SOM-based Lorenz curve analysis allows quantification of unequal spatial distribution

If a genomic activity is spatially constrained within the nucleus (e.g. it appears only in a particular compartment), the loci displaying that activity should be non-uniformly distributed on the trained SOM lattice. We can quantify the degree of non-uniformity in SOM distribution by performing Lorenz curve analysis. Lorenz curves represent the cumulative proportional distributions of a resource over a population [31]; they are often used to represent income or wealth inequality over a population. A Lorenz curve displays the cumulative proportion of the population that has a given cumulative share of the resource, after ordering the population from lowest share to highest. Statistics such as the Gini coefficient quantify the degree of resource-sharing inequality in a given population by comparing the area under the observed Lorenz curve to a distribution representing equal resource distribution (i.e. defined by the diagonal) [33]. In our usage, we make Lorenz curves based on the distribution of a given genomic activity over the SOM lattice. Since genomic loci are not equally distributed on the lattice, we compare the area under the observed Lorenz curve to a curve defined by the distribution of random loci assignment (**Figure 5A**). The resulting modified Gini coefficient measures the degree to which a given activity is non-uniformly distributed over the SOM, and ranges from zero (representing a uniform distribution) to one (representing a highly non-uniform distribution).

**Figure 5:**
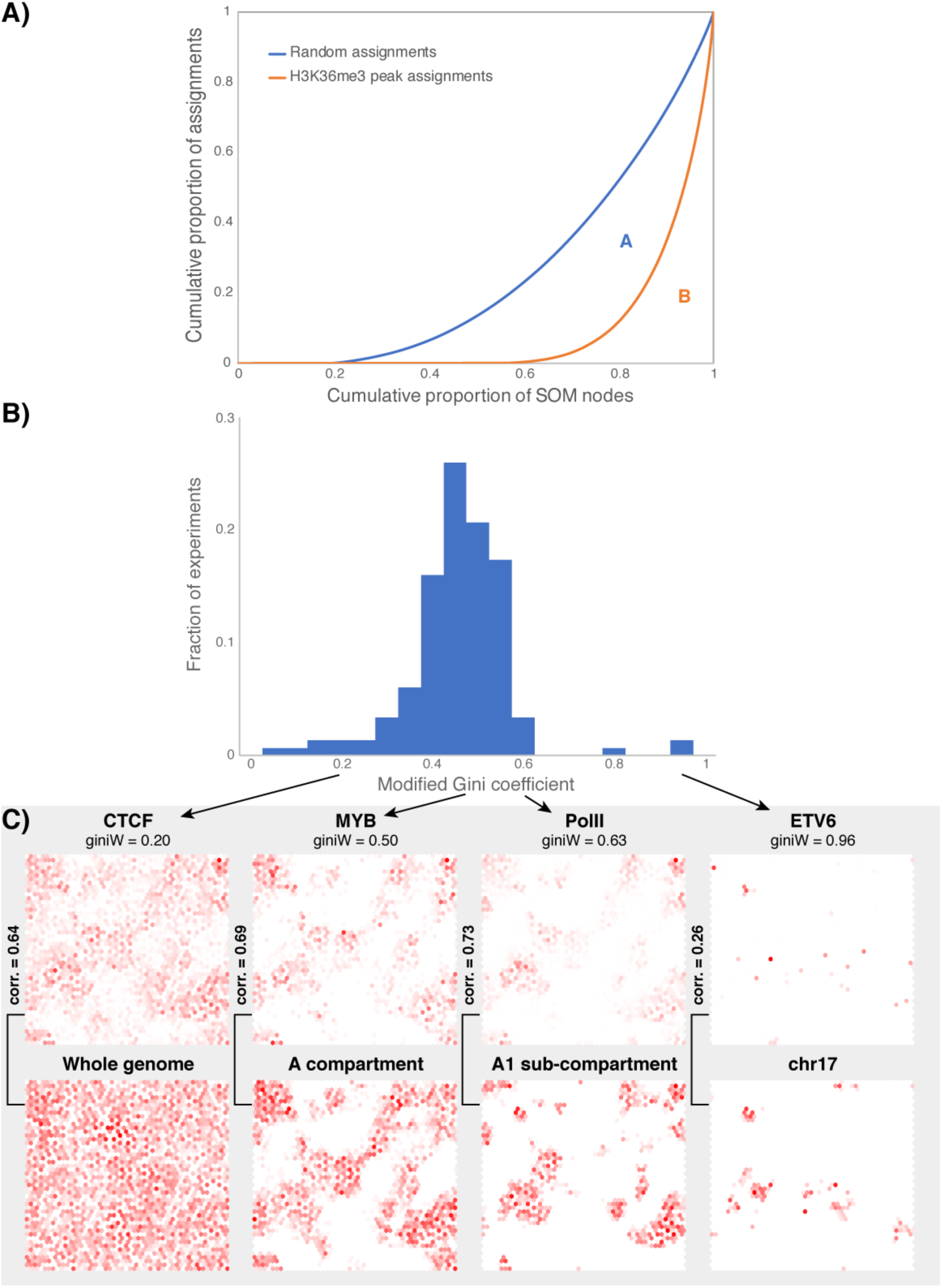
**A)** Illustration of Lorenz curves for the SOM projections of H3K36me3 peaks (orange curve) and random loci (blue curve). The modified Gini coefficient is calculated as A/(A+B), where the areas A & B are calculated as displayed on the graph. **B)** Distribution of modified Gini coefficients calculated by projecting peaks from 150 transcription factor ChIP-seq experiments onto the trained GM12878 SOM. See **Supp. Table 1** for values. **C)** Projections of selected transcription factor ChIP-seq experiments onto the SOM, and comparison with relevant projections of compartments or chromosomes.

We used our trained GM12878 SOM and Lorenz curve analysis to assess the spatial distributions of peaks from 150 transcription factor ChIP-seq experiments performed by the ENCODE consortium. Our results show that most of the experiments display intermediate range Gini coefficients, with a median score of 0.49 (**Figure 5B, Supp. Table 1**). Most such intermediate scores can be explained by a general association between the relevant transcription factor’s peaks and the A compartment. For example, the distribution of MYB’s peaks on the SOM are highly correlated with the distribution of A compartment loci on the SOM (**Figure 5C**). Interestingly, peaks from PolII ChIP-seq experiments have higher than average Gini coefficients, suggesting a more constrained spatial localization, and their SOM distribution is more tightly correlated with that of the A1 sub-compartment (**Figure 5C**). This result is consistent with previous observations that A1 represents a sub-compartment with higher regulatory activity within the nucleus [27].

Conversely, peaks from CTCF and cohesin (SMC3 & RAD21) display lower Gini coefficients, reflecting their more uniform spread throughout the SOM (**Figure 5C**). These results are consistent with the appearance of CTCF and cohesin peaks at the boundaries between A and B compartment TADs, which would therefore appear to have no spatial localization within the SOM (at least at the 250kbp resolution used in this study). Finally, outlier high Gini scores can sometimes be explained by high association with a particular chromosome, as opposed to more general forms of compartmentalization. For example, some ETV6 ChIP-seq experiments display high Gini scores, but this is explained by highly disproportionate numbers of peaks appearing on certain chromosomes (chr3, chr17, and chr20). While this would be a valid form of spatial localization within the nucleus, it may also be due to technical artefacts in the ChIP-seq experiments or peak-finding analyses. We note that other ETV6 experiments display intermediate Gini scores (**Supp. Table 1**) and do not display disproportionate associations with specific chromosomes.

We also performed Lorenz curve analysis on the 16 DNase-seq and histone modification ChIP-seq experiments (**Supp. Table 2**), and on annotations for 41 chromatin states produced by the IDEAS platform [30] (**Supp. Table 3, Figure S1**). The results of these analyses are consistent with the discussion of the transcription factor ChIP-seq experiments; higher Gini scores generally correspond to more specific localization in the A compartments.

## DISCUSSION

We have introduced a new SOM-based approach for visualizing chromatin organization in 2D. Since the trained SOM lattice contains discrete entities (nodes) that each encapsulate a set of genomic loci, it is straightforward to project any genomic activity onto the SOM, even if the loci displaying that activity are measured at a different resolution to the SOM training set. We can thereby easily visualize how genomic activities are distributed over the 2D space defined by the SOM lattice, which is implicitly related to the distribution of that activity within the nucleus. We have demonstrated that it is easy to measure relationships between the SOM distributions defined by distinct genomic activities. We have also shown that the degree of non-uniformity displayed by a genomic activity on the SOM (and hence within the nucleus) can be conveniently measured using Lorenz curve analyses.

One disadvantage of our approach is that the time taken to train a SOM becomes computationally prohibitive with large numbers of genomic loci (i.e. smaller genomic intervals produced from higher resolution Hi-C data). However, SOM training only has to be performed once per Hi-C dataset. Projecting a genomic activity onto a trained SOM is not a costly operation, as it consists of merely comparing the genomic coordinates that display the activity to the genomic loci encapsulated in each of the SOM’s nodes.

In summary, we have demonstrated that our approach to dimensionality reduction of chromatin interaction data enables a unique way to integrate large numbers of epigenomic datasets in the context of chromatin organization.

## ACKNOWLEDGEMENTS

This material is based upon work supported by the National Science Foundation under ABI Innovation Grant No. DBI1564466 (to SM). Any opinions, findings, and conclusions or recommendations expressed in this material are those of the author(s) and do not necessarily reflect the views of the National Science Foundation. TK was also supported by an Erikson Discover Grant and the Edward B Nelson Undergraduate Research Fund, both from Penn State University.

## AVAILABILITY

Open source Java code (MIT license), an executable JAR file, and scripts for reproducing this manuscript’s analyses are available from https://github.com/seqcode/chromosom.

## CONTRIBUTIONS

SM designed and oversaw the study. TK wrote the ChromoSOM code and performed analyses of data projected to the SOM, with contributions from SM. LR wrote scripts for Hi-C data processing and performed Hi-C normalization. SM and TK wrote the manuscript. All authors approved the final manuscript.

## SUPPLEMENTAL MATERIALS

**Figure S1:**
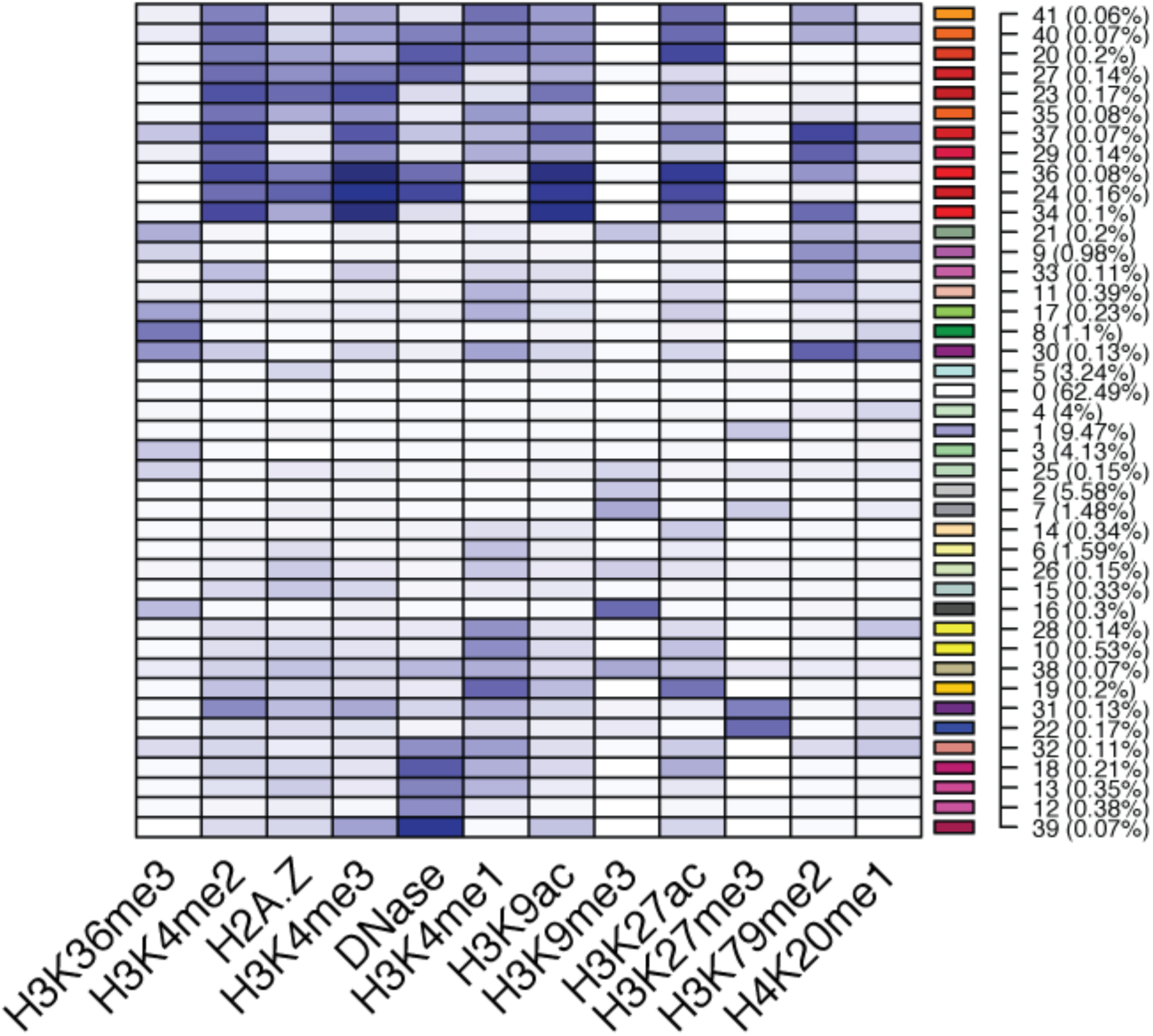
Chromatin mark enrichment patterns for IDEAS states. Reproduced from [30], which is the source of the state annotations used in this study. Numbers in brackets represent the percentage of the genome covered by each state. This figure should be used as a guide to interpret the state numbers defined in **Supp. Table 3.**

**Supplemental Table 1:**
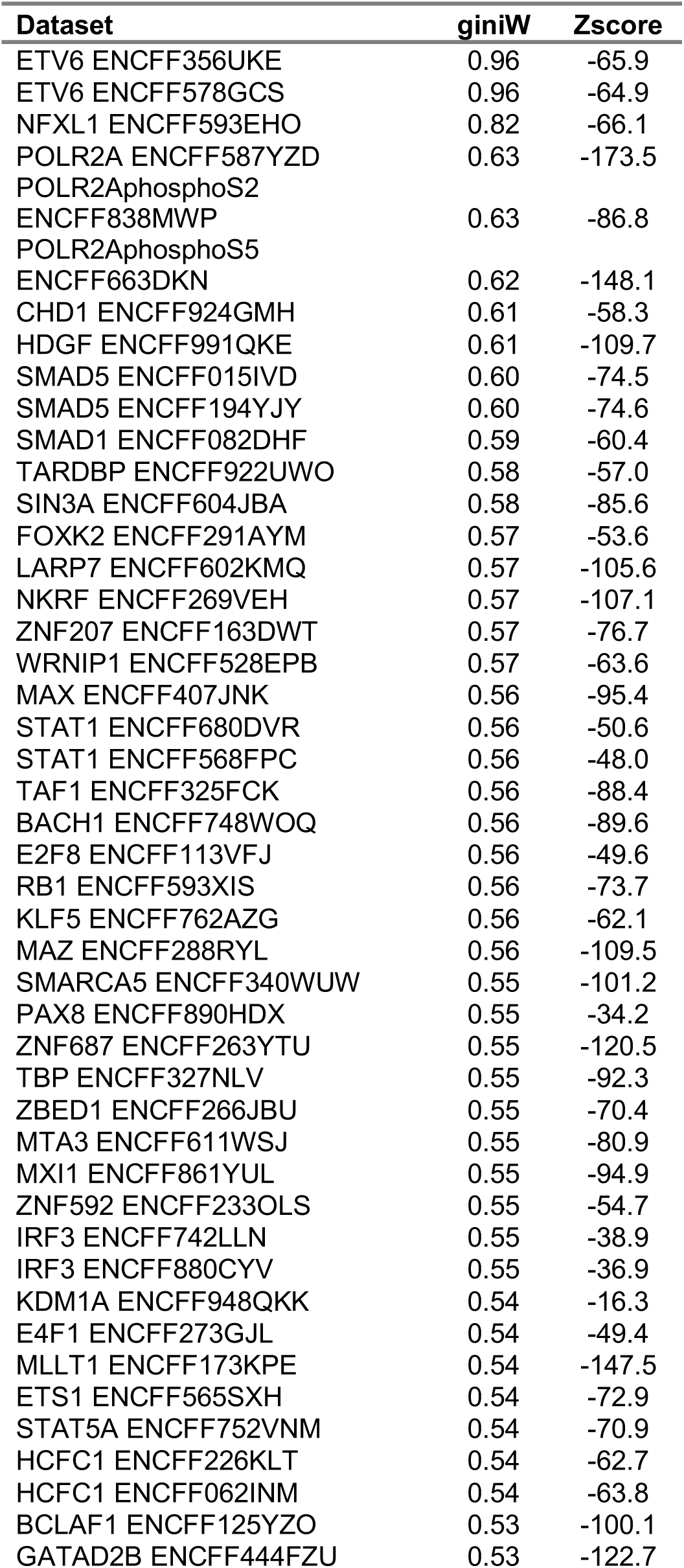
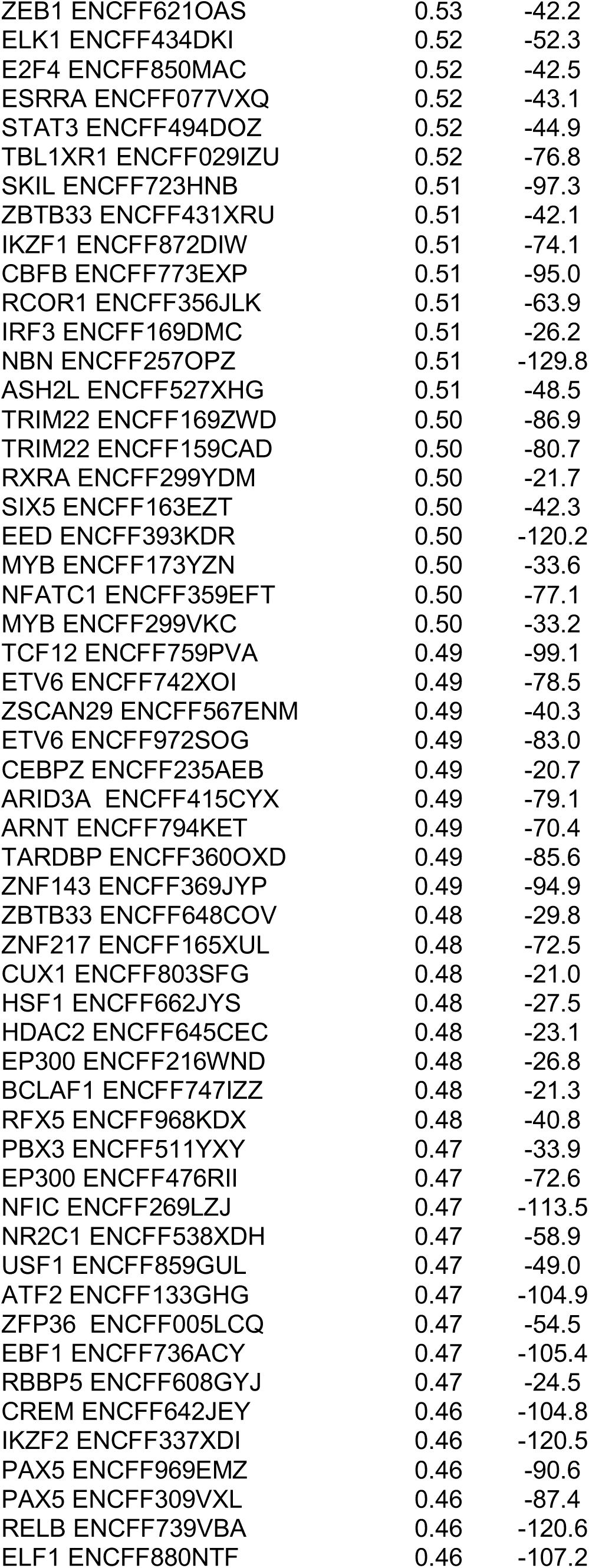
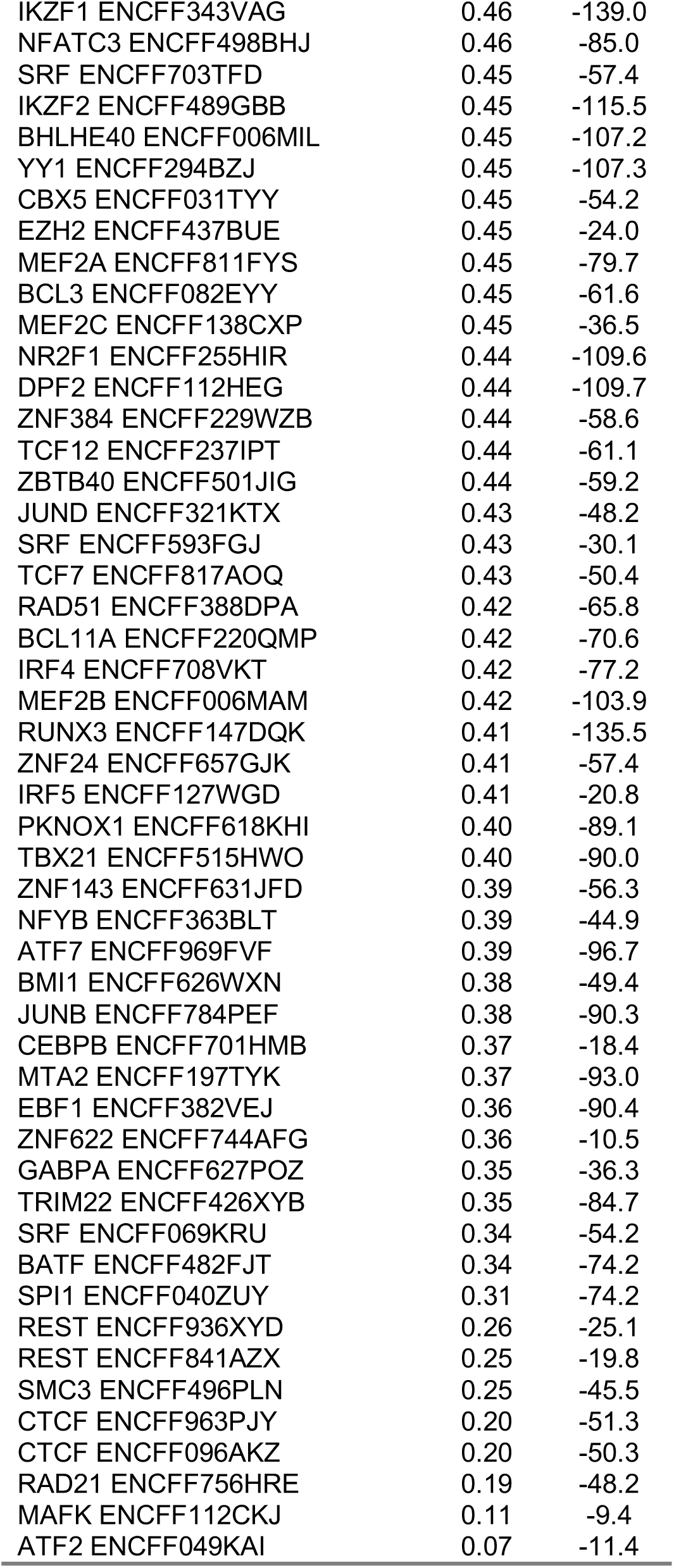
Modified Gini coefficients (giniW) calculated for the SOM projections of peaks from 150 GM12878 transcription factor ChIP-seq experiments.

**Supplemental Table 2:** Modified Gini coefficients (giniW) calculated for the SOM projections of peaks from 16 GM12878 DNase-seq and histone mark ChIP-seq experiments.

| **Dataset** | **giniW** | **Zscore** |
| --- | --- | --- |
| H3K36me3 ENCFF479XLN | 0.65 | -262.9 |
| H4K20me1 ENCFF308WNH | 0.62 | -68.9 |
| H3K79me2 ENCFF357HZM | 0.62 | -265.9 |
| H3K9ac ENCFF052MHA | 0.55 | -143.9 |
| H3K27ac ENCFF816AHV | 0.52 | -166.3 |
| H3K27me3 ENCFF851UKZ | 0.51 | -108.4 |
| H3K4me3 ENCFF354MGT | 0.51 | -128.4 |
| H3K4me3 ENCFF795URC | 0.50 | -151.7 |
| H3K4me1 ENCFF921LKB | 0.50 | -194.7 |
| H3K4me3 ENCFF295GNH | 0.49 | -129.4 |
| H3K4me2 ENCFF983SMS | 0.45 | -177.4 |
| H3K27me3 ENCFF247VUO | 0.43 | -91.2 |
| H2A.Z ENCFF584NAD | 0.43 | -144.1 |
| DNase-seq ENCFF804BNU | 0.37 | -320.8 |
| DNase-seq ENCFF097LEF | 0.33 | -100.5 |
| DNase-seq ENCFF273MVV | 0.25 | -90.5 |

**Supplemental Table 3:** Modified Gini coefficients (giniW) calculated for the SOM projections of peaks from IDEAS chromatin state annotation of the genome using GM12878 chromatin modifications.

| Dataset | giniW | Zscore |
| --- | --- | --- |
| state41 | 0.91 | -118.8 |
| state25 | 0.90 | -173.3 |
| state29 | 0.87 | -38.4 |
| state40 | 0.81 | -102.5 |
| state19 | 0.80 | -68.7 |
| state5 | 0.78 | -226.7 |
| state21 | 0.70 | -94.0 |
| state20 | 0.70 | -69.0 |
| state9 | 0.69 | -181.6 |
| state37 | 0.66 | -92.4 |
| state30 | 0.64 | -145.1 |
| state36 | 0.62 | -60.3 |
| state17 | 0.62 | -48.5 |
| state27 | 0.61 | -39.5 |
| state32 | 0.60 | -150.6 |
| state8 | 0.59 | -236.3 |
| state16 | 0.58 | -129.7 |
| state6 | 0.58 | -148.7 |
| state34 | 0.57 | -74.3 |
| state3 | 0.55 | -370.0 |
| state33 | 0.53 | -170.7 |
| state38 | 0.53 | -45.0 |
| state1 | 0.50 | -202.4 |
| state39 | 0.48 | -91.8 |
| state28 | 0.48 | -194.7 |
| state4 | 0.46 | -93.0 |
| state26 | 0.44 | -62.8 |
| state35 | 0.42 | -84.6 |
| state22 | 0.42 | -31.7 |
| state13 | 0.41 | -89.0 |
| state7 | 0.39 | -42.5 |
| state31 | 0.32 | -14.9 |
| state12 | 0.27 | -67.9 |
| state15 | 0.23 | -30.1 |
| state2 | 0.18 | -134.8 |

